# Real-time assessment of the impacts of polystyrene and silver nanoparticles on hatching process and early-stage development of *Artemia* using a microfluidic platform

**DOI:** 10.1101/2023.08.16.553636

**Authors:** Preyojon Dey, Terence M. Bradley, Alicia Boymelgreen

## Abstract

The development of real-time in-situ monitoring techniques is key to advancing a mechanistic understanding of the impacts of marine pollution, which is challenging to acquire through traditional end-point toxicity testing. We investigated the impacts of different nanopollutants on the hatching process and early-stage development of marine organisms, a vulnerable life stage, by observing oxygen consumption in real-time and morphological changes at regular intervals using a microfluidic platform. Here, two common and distinct nanoparticle (NP) types - polystyrene (PS) nanoplastic and silver (Ag) nanometal, were examined to assess and compare impacts on the hatching process and nauplius stage (first larval stage) of *Artemia*, a widely used zooplankton model in ecotoxicological studies. The study was conducted over a wide range of doses that are relevant to different environmental conditions, ranging from 0-1 mg/L, over a period of 24 hours. The hatching process of *Artemia* is comprised of four distinct stages which can be differentiated by metabolism and morphology: hydration, differentiation, emergence, and hatching. During hatching, NP exposure altered the time needed for the resumption of dormant *Artemia* cysts (hydration duration) at the lowest dose, dramatically prolonged the differentiation stage, and slowed embryo emergence from the cysts. The remaining time for the hatching stage during the experimental timeframe was also shortened. Overall, the presence of NPs led to increased oxygen consumption in multiple stages of the hatching process. Hatchability increased significantly with NP concentration although mortality showed an inverse pattern. This may be attributed to the increased aggregation of NPs in saltwater with increasing concentration which limits bioavailability during hatching but may be more readily consumed post-hatch. Ag NPs had a greater effect on hatching and mortality in comparison to PS NPs. A significant impact of NPs on swimming speed was observed, with a decrease observed in the presence of PS NPs and an increase observed in the presence of Ag NPs.

**Graphical abstract:** 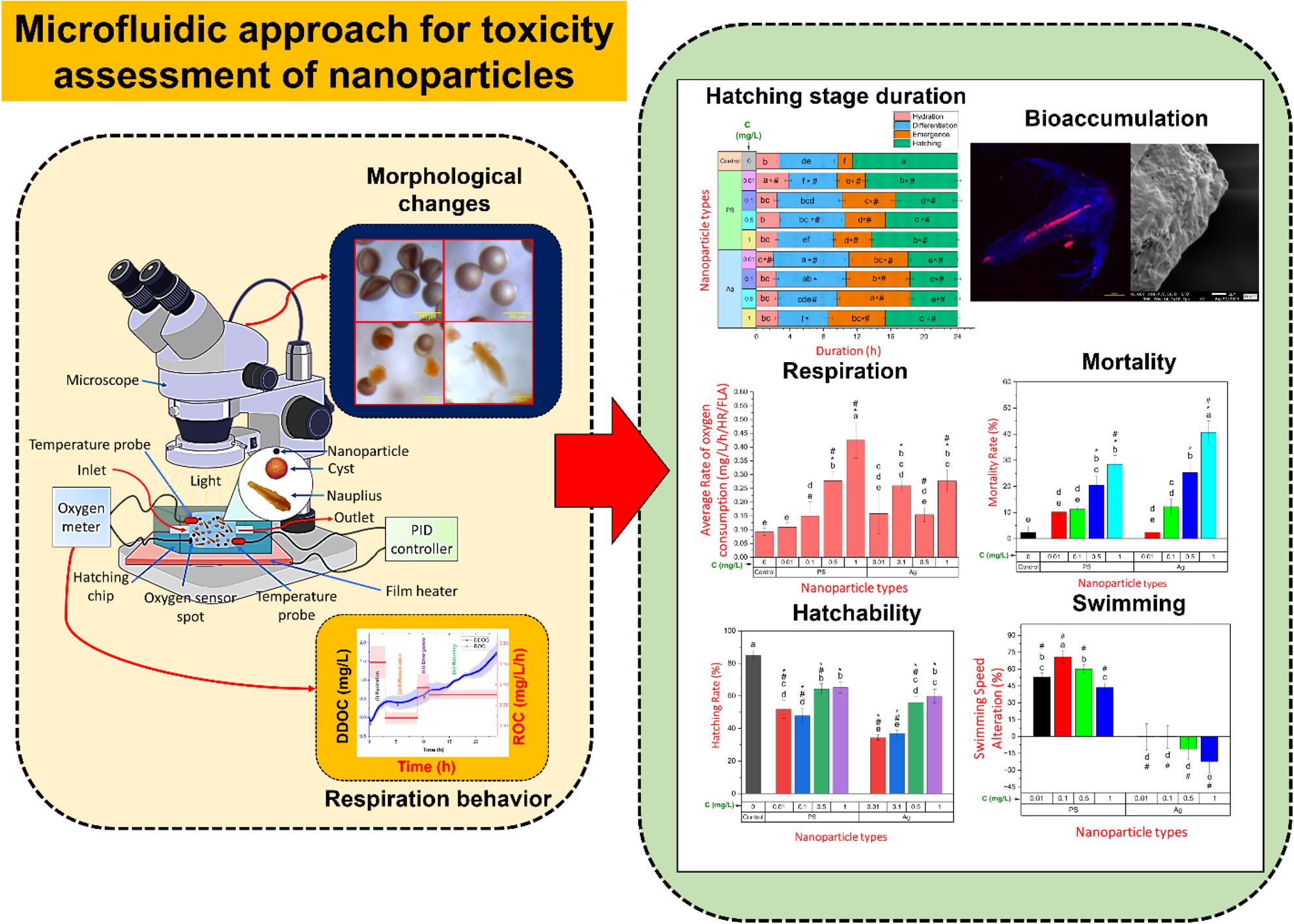

**Highlights:**

- Utilization of oxygen sensor integrated microfluidic chip and microscopy for ecotoxicological study.
- Bioaccumulation of NPs affected hatching stages and respiration leading to inhibition of hatchability, with greater toxicity of silver NPs.
- NPs caused significant mortality and alteration in swimming performance.

## 1. Introduction

Rivers, runoff, and direct discharges carry a wide range of contaminants from anthropogenic activities into the ocean. Plastic is a major marine pollutant, accounting for approximately 80-85% of the total marine litter ^1–3^. Every year, almost 8 million tons of plastic garbage are discharged into the ocean ^2^. This massive volume of plastic waste can disintegrate into micro and nanoparticles (MNPs) by a variety of processes including mechanical degradation, photo-degradation, and hydrolysis ^4,5^. Nanotechnology has increased the demand for and production of metallic nanoparticles (NPs), which are now emerging as a new form of marine pollution ^6,7^. Various studies have been conducted on the effects of these small-sized marine pollutants primarily focused on microparticles (MPs) ^8–11^. Evidence suggests NPs can cause more harm to marine life ^12^ due to size-dependent properties (smaller size and weight, higher surface area, etc.), which allow them to spread more readily ^13,14^, be misinterpreted by marine animals for food ^15,16^ and be absorbed and promptly release toxins ^17,18^. In particular, the reproductive and early stages of marine species can be severely impacted by NPs due to the vulnerability of this phase of development ^19,20^, potentially impacting abundance and distribution, and the marine ecosystem ^21,22^. The heterogeneity of nanopollutants in the marine environment can also modify these effects ^23,24^ supporting the need for comparative analyses of the impacts of different types of nanoparticles particularly at environmentally relevant concentrations.

This study employed two types of model nanoparticles: polystyrene (PS) and silver (Ag). PS accounts for 23 million tons of global plastic production per year ^25^. Roughly 3% of the total produced plastics end up in the ocean ^26^, with polystyrene (PS) being identified as a prevalent polymer found in marine debris ^27–29^. The plastic pollutants exhibit a range of sizes, with a majority of 70-80% classified as macroplastics ^30^, measuring 5-100 mm ^31^. While 15-30% of the plastic waste is micro/nanoplastics, the macroplastics also undergo fragmentation over time, leading to the formation of secondary microplastics and nanoplastics ^30^. PS nanoplastics are utilized in packaging and nanomedicine applications as well. Ag NPs are one of the most commonly used metallic nanoparticles in medical applications ^32^, biosensors ^33^, nano-enabled consumer electronics ^34^, textiles and building facades, which may emit Ag NPs that end up in the water ^35,36^ and are highly toxic ^37^. It is important to note that the behavior of MNPs in fresh and saltwater differs significantly. In saltwater, MNPs tend to agglomerate owing to the loss of electrostatic repulsion in the presence of ions ^38^. This phenomenon has the potential to impact the bioavailability of MNPs and consequently can affect aquatic organisms in a manner distinct from that observed in freshwater environments ^39–41^.

Previously, PS and Ag NPs have been studied on *Brachionus plicatilisi* ^42^,and *Oryzias latipes* ^43^, revealing that both types of NPs are toxic. Here, a zooplankton species, commonly used as live feed in marine finfish larviculture due to its small size and rich nutrient content ^44,45^, *Artemia franciscana*, was used to examine the toxic effects of these two NPs on both the hatching process and nauplius stage. Previously, this species has been widely used as an aquatic model animal in ecological, ecotoxicological, genetic, biochemical, and physiological studies ^33–35^. Previous comparative studies on the impacts of different NPs on *Artemia* focused primarily on the larval or adult stages ^46–52^ with few studies evaluating the impacts of different NPs on hatching rate (HR) ^53–56^. These studies demonstrated a significant decrease in hatching rate in the presence of NPs, stimulating investigation into the mechanisms underlying the inhibition of hatching.

This study provides a comparative analysis of the impacts of two types of nanoparticles, PS and Ag on the hatching and early stage of *Artemia* under identical environmental conditions. While the effects of NPs on hatching rate have been examined using conventional endpoint toxicity assessment along with survival and mobility, this study expands the understanding of these impacts beyond the endpoint to include real-time information on the individual stages of the hatching process and early-stage development. This was achieved through the development of a microfluidic platform integrated with an optical oxygen sensor that tracks respiration of *Artemia* in real-time throughout hatching. Physical changes in *Artemia* during hatching were recorded by a microscope and correlated to the sensor data for estimation of the effect on the hatching process ^57^. Automatic handling and counting of test animals on a microfluidic platform enhanced accuracy during hatchability estimation ^57^. Previously, micro and milli-fluidic platforms have been utilized in several studies on live animals ^58–60^, including zebrafish ^61–63^ and various respirometry techniques were used to quantify oxygen consumption of different marine species, such as *Perna viridis* ^64^, *Dendropoma cristatum* ^65^ and *Palaemon adspersus* ^66^ exposed to different marine contaminants. In a previous study by our group ^57^, an on-chip optical oxygen sensor was utilized to monitor oxygen consumption of *Artemia* during hatching under different temperatures and salinities, and influence of temperature and salinity on the duration of hatching stages, metabolic rates, and hatchability could be evaluated on-chip. Due to its compact size, the microfluidic platform can prevent sensor noise ^67–72^ and provide consistent hatching temperature via an off-chip controller. It is anticipated that the present approach could be employed to shed light on the effects of different marine nano-pollutants on the hatching and early development of a variety of other marine species.

## 2. Experimental

### 2.1 Nanoparticles and their characterization

Red fluorescent PS NPs of 50 nm nominal size dissolved in an aqueous solution at a concentration of 1% solids (w/v) were purchased from Thermo Scientific Chemicals. Citrate-capped Ag NPs of nominal size 40 nm diluted in an aqueous buffer containing sodium citrate as a stabilizer at a concentration of 0.02 mg/ml were purchased from the same vendor. As-received NP solutions were drop cast onto a double-sided carbon tape adhered to a glass coverslip and air dried overnight prior to imaging with scanning electron microscopy (SEM) (JEOL FS-100). SEM images indicated that PS NPs were consistently spherical in shape (Figure 1A), whereas Ag NPs had an irregular shape (Figure 1B). PS and Ag NPs had a Z-average size of 50.43 ± 0.24 nm and 45.83 ± 0.13 nm, a polydispersity index (PDI) of 0.02 ± 0.01 and 0.23 ± 0.00, and a Zeta potential of −23.8 ± 0.96 mV and −34.07 ± 1.86 mV, respectively, as determined by dynamic light scattering (DLS) (Nano ZS, Malvern instruments) (results are expressed as mean ± standard deviation). PS and Ag NPs were suspended in varying amounts with artificial saltwater (ASW) of 25 parts per thousand (ppt) salinity using a vortex mixer (Vortex genie 2, Scientific Industries) to make NPs solutions with concentrations of 0.01, 0.1, 0.5, and 1 mg/L. The concentration of marine pollutants, including plastic, can vary across different locations ^73,74^. In certain regions, concentrations of plastic as high as 1.26 mg/L have been reported ^75^. ASW was prepared by adding commercially available sea salt (Fluval Sea, pH: 8.1-8.2) to deionized water (DIW) (prepared using Barnstead Smart2Pure Water Purification System, Thermo Scientific). Thorough mixing of the ASW solution permits homogeneous distribution of the dissolved oxygen and salt. The NP solutions were kept at 25⁰C for 24 h. The behavior of nanoparticles in saltwater in different concentrations was tested by using DLS after 1 and 24 h. Before testing, the ASW solution was filtered (MS sterile syringe filter, pore size= 0.22 µm), and mixed with NPs of varying concentrations. The Z-average size and PDI of the NPs in the ASW at each concentration were determined by DLS (10 runs of 10 sec for each sample) after 1 and 24 hours at 25°C from experiments performed in triplicate. Additionally, photographs were captured at the bottom of the tubes to assess the likelihood of sedimentation after 24 h.

**Figure 1.**
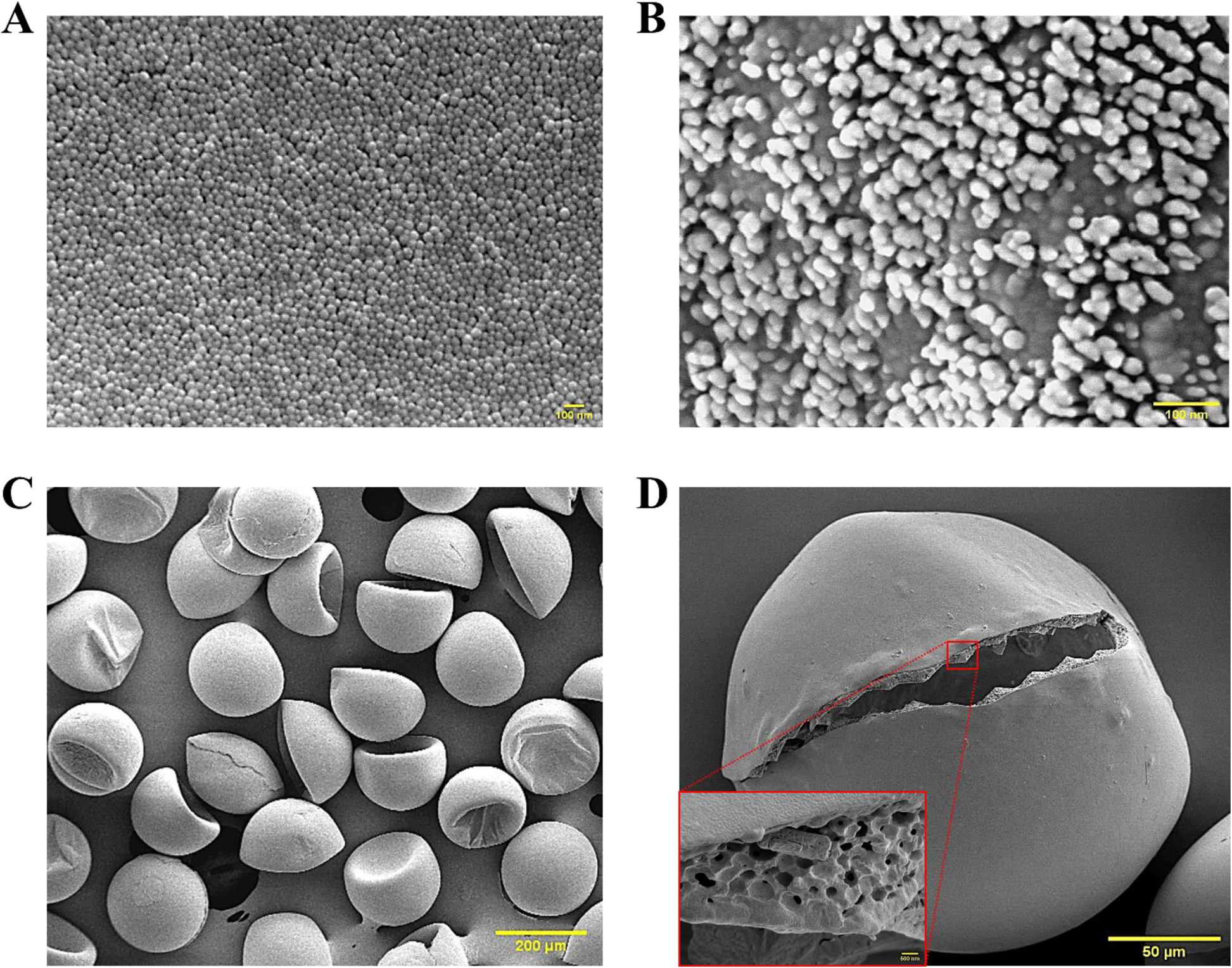
Scanning electron microscopy (SEM) images of- (A) Polystyrene (PS) nanoparticles (scale bar= 100 nm). (B) Silver (Ag) nanoparticles (scale bar= 100 nm). (C) *Artemia* cysts before hatching (as received) (scale bar= 200 µm). (D) Outer surface lamella and inner surface 3D porous structure (inset, scale bar= 500 nm) of *Artemia* cysts (scale bar= 50 µm).

### 2.2 *Artemia* cysts and hatching

*Artemia* cysts were procured from Brine Shrimp Direct and stored at 4⁰C until one day before the hatching experiments, when they were moved to room temperature to allow for gradual temperature adaptation. The as-received cysts were mixed with ASW (control) or ASW with varying concentrations of NPs at 5 g/L cyst concentration and hatched for 24 hours using a microfluidic platform. The design, fabrication, and operation of the microfluidic platform were discussed in detail in our previous work ^57^. Briefly, a polydimethylsiloxane (PDMS) hatching chip, with a cylindrical enclosed chamber with inlet and outlet was utilized. Chip fabrication involved PDMS casting in a 3D-printed mold (fabricated by using Asiga MAX X43 3D printer and SuperCAST V3 resin). The chip was connected to a thermocouple probe (TC 10K, QINGDAO) and placed on a proportional–integral– derivative (PID) controlled film heater (PI film heater-24 V, Icstation) to maintain a uniform temperature for hatching (25⁰C). The selection of a temperature of 25⁰C and a salinity of 25 ppt for the ASW was based on the findings of our previous study, which demonstrated that these conditions yielded optimal hatching performance ^57^. An oxygen sensor spot (OXSP5, Pyroscience) was placed on the wall of the hatching chip chamber and connected to an oxygen meter (FireSting®-O_2_, Pyroscience) through an optical fiber. With the oxygen-dependent luminescence emission from oxygen-sensitive spot, the oxygen meter can measure dissolved oxygen concentration of the hatching media within the chip every second. For hatching experiments, cyst solutions in ASW or NP-spiked ASW were inserted into the hatching chip, and its inlet and outlet were sealed with binder clips to avoid air infiltration. The entire hatching setup was placed under a digital stereo optical microscope (SE400-Z, Amscope) equipped with a digital camera (MD500, Amscope) to capture photomicrographs of the hatching process every 5 minutes. Due to the stimulatory effect of light on hatching ^76,77^, a light-emitting diode (LED) light (1W, Amscope) was continuously on throughout the hatching process. At least three hatching experiments were conducted at each NP concentration. Supplementary Figure 1A depicts a schematic diagram of the experimental setup utilized for hatching, and Supplementary Figure 1B illustrates the computer aided design (CAD) model of the hatching chip.

Scanning electron microscopy (SEM) (JEOL FS-100) was utilized to study the morphology of the *Artemia* cysts. To obtain SEM images, the nonconductive cysts underwent a gold-coating process. SEM images reveal that as-received cysts had a cup-shaped structure (Figure 1C) with a smooth outer surface lamella and the inner cyst wall had a 3D porous structure with pores ranging from 160-900 nm (Figure 1D). Due to this unique structure, NPs present in the environment can be entrapped into these pores ^78,79^ which may affect the hatching process. NPs tend to agglomerate in saltwater, resulting in the formation of aggregates that are substantially larger than the initial NP size (elaborated in section 3.1). To test whether these large particles are covering the cysts, or diffusing into the 3D porous structure, some *Artemia* cysts were imaged under SEM, which were hatched in the presence of different sized PS (50 nm, 2 µm and 10 µm) and Ag NPs (40 nm and 100 nm) at 1 mg/L concentration. To obtain SEM images of the cysts, it was necessary to desiccate them (done by air drying at room temperature) prior to imaging. For this purpose, the cysts were not hatched in ASW but DIW, as the former results in the formation of sizable salt particles that obscure the morphology of the cysts [Supplementary Figure 2 (A)]. The use of DIW rather than ASW resulted in hatching taking longer than 24 h ^57^, however, during the 24 h time period, many cysts were cracked open (due to increased internal turgor pressure, described in detail in section 3.2) and were visualized under SEM. Various sizes of nanoparticles were employed to simulate the agglomeration of nanoparticles in saltwater during the hatching process. Energy dispersive X-ray spectrometry (EDS) (JEOL FS-100) was used to assess the distribution of Ag NPs on the cysts. The results are presented in Figure 4 and Supplementary Figure 2.

### 2.3 Duration of different stages, and rate of oxygen consumption

The on-chip oxygen sensor monitored the dissolved oxygen concentration (DOC) of the water within the chip. As *Artemia* cysts undergo the process of hatching, they consume oxygen, which causes the concentration of dissolved oxygen within the chip to decrease over time. This depletion in dissolved oxygen concentration (DDOC) at one second intervals was calculated by subtracting the DOC of every given second from the DOC of the previous second. Supplementary Figure 3 illustrates DDOC vs hatching time for *Artemia* under various NPs treatments as well as for a blank experiment (ASW with no cysts). Optical photomicrographs of the morphological alterations of the hatching cysts and the subsequent change in the overall trend of DDOC were utilized to determine the duration of the various stages of hatching.

Oxygen consumption at any hatching stage or DDOC at that stage (difference in DOC at the beginning and end of any stage) is normalized based on duration of that particular stage, since the absolute value and duration of various stages of hatching vary with concentrations and types of NPs (discussed later in section 3.3). The measurement of oxygen consumption represents the cumulative value of consumption by all viable cysts present throughout the initial three stages of the hatching process (hydration, differentiation and emergence), as well as the number of successfully hatched *Artemia* and those surviving post hatch. Accordingly, the average rate of oxygen consumption (aROC) of *Artemia* during the hatching process is normalized by the duration and hatching rate and divided into two distinct phases: A) the average rate of oxygen consumption during any of the first three stages or pre-hatching rate of oxygen consumption (aROC_1-3_), and B) the average rate of oxygen consumption during the hatching stage or post-hatching rate of oxygen consumption (aROC_4_). aROC_1-3_ is normalized based on the duration of the any of the first three stages multiplied by the hatching rate assuming the proportion of viable cysts during this stage is equivalent to the hatching rate. This is expressed by the following formula:

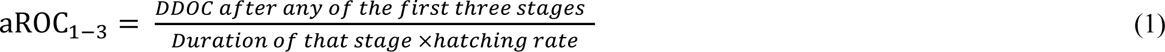

where the hatching rate is calculated according to the following formula:

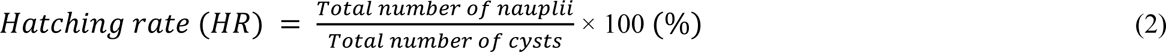

The methodology for determining the hatching rate is described in section 2.5.

In contrast, aROC_4_ is normalized based on the duration of the final stage of hatching (hatching stage), hatching rate and fraction of live *Artemia* (FLA). This is expressed by the following formula:

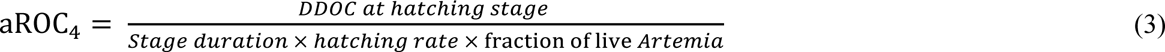

where the fraction of live *Artemia* (FLA) is calculated according to

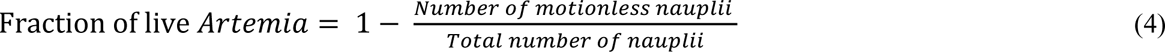

### 2.4 Mortality and swimming speed alteration tests

Mortality rate and swimming speed of newly hatched *Artemia* were evaluated as conventional endpoint toxicity tests for NPs, as reported in previous studies ^80,81^. Twenty-four hours after initiation of the experiment (mixing of cysts and ASW), cysts, and nauplii were transferred from the hatching chip to a shallow depth (1.2 mm) counting chip using a continuous flow of ASW from the syringe pump (100 µL/min) at the hatching chip inlet. The shallow depth forces cysts and nauplii into a monolayer facilitating counting while still allowing free movement. The chip outlet has micropillars that permit water flow but prevent cysts and nauplii from escaping (Supplementary Figure 1C). The transferred cysts and nauplii in the counting chip were manually agitated for 15 sec, and then imaged with the stereo optical microscope. Nauplii that were not motile were considered dead and the mortality rate was calculated using the following formula:

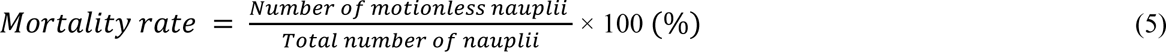

For swimming speed experiments, approximately 18-20 nauplii were transferred back to the hatching chip using the backflow of ASW from the counting chip outlet and the chip was filled with ASW (no NP). Nauplii in the hatching chip were held in the dark for five minutes to allow them to acclimatize, uniformly distribute and achieve a consistent speed ^82,83^. After five minutes of darkness, a one-minute video of the swimming nauplii was captured under light conditions with the same digital camera equipped stereo optical microscope. The nauplii swimming path was tracked in ImageJ until individuals exited the frame, and the average swimming speeds (S) were then calculated. Only the tracking paths of nauplii that remained in at least ten frames were analyzed. The mortality and swimming speed of nauplii exposed to each NP concentration were measured in at least three distinct experiments, and in each experiment, the swimming speed of at least eight nauplii was measured. The swimming speed alteration (SSA) of the nauplii exposed to NPs was calculated with the following formula used in the literature ^80^:

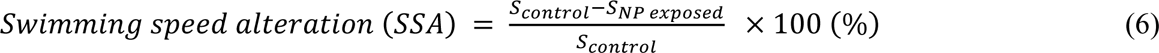

### 2.5 Hatching rate (HR) estimation

The nauplii and cysts (hatched or unhatched) in the counting chip were euthanized through exposure to a continuous flow of 30% methanol solution in ASW using a syringe pump. The 30% methanol solution euthanizes *Artemia* nauplii in a few minutes. Post euthanasia, cysts and nauplii were photographed using a digital camera (D3100, Nikon), and the photos were analyzed using ImageJ to count the number of nauplii and cysts based on their circularity (if circularity ≥ 0.9, it was considered a cyst, otherwise a nauplius). The total number of nauplii was calculated by adding the number of transferred nauplii for swimming tests to this number. Further details on the automated counting approach based on the image are available in Ref ^57^. The hatching rate of the *Artemia* in different NP exposure conditions was measured by Formula 2, described in section 2.3. The hatching rate at each NP condition was measured from at least three experiments.

### 2.6 Uptake of nanoparticles

The uptake of both NPs by *Artemia* hatched under various nanoparticle treatments was measured with bright field images with an optical microscope (Axioscope 5, Zeiss) and a digital camera (Axiocam 208 color). A spinning disc confocal fluorescence microscope (Nikon Eclipse Ti2-E) was employed to also image *Artemia* nauplii that were exposed to fluorescent PS NPs and the fluorescent intensity of the NPs were measured by using ImageJ program (Supplementary Figure 4). All photos were captured from immobilized *Artemia* collected from the hatching chip after 24 hours of exposure to NPs and placed on glass slides.

### 2.7 Statistical analysis

For all statistical analyses, OriginPro (version 2023b, OriginLab) was utilized. The data are displayed as the mean ± standard deviation. A one-way analysis of variance (ANOVA) followed by the Tukey post-hoc test was performed to determine the significance of the differences between treatments and the control group. Prior to performing ANOVA tests, the data were tested for normality using a normal quantile-quantile plot with a confidence level of 95%. In this study, two separate correlation analyses were employed, utilizing the Pearson correlation coefficient, due to the inability to ascertain the effects of hatching in the presence of NPs on the post-hatching stage. The first analysis examined the parameters investigated up to the hatching of *Artemia*, including the duration of all stages, aROC, and hatching rate. The second analysis focused on the post-hatching stage, examining the duration of only the hatching stage, aROC at hatching stage, mortality rate, and swimming speed alteration (at least 27 NP treatment conditions were considered for analysis of each category of data, see Supplementary Figure 5). If *p*<0.05, the results were considered significant in all statistical analyses.

## 3. Results and Discussion

### 3.1 Aggregation behavior of PS and Ag nanoparticles in saltwater

High ionic strength reduces the thickness of the electric double layer (EDL), and decreases electrostatic repulsion, leading nanoparticles in saltwater to interact, agglomerate, and settle relative to freshwater ^84,85^. Higher temperature (25 ⁰C) can also promote NP aggregation ^86^. This agglomeration is expected to affect the uptake of NPs since the effective radius is altered ^87,88^. Figure 2 depicts the hydrodynamic size (Z-average) and polydispersity index (PDI) of different concentrations of PS and Ag NP solutions in ASW after 1 and 24 hours. PS NPs aggregated more than Ag NPs to form microscale aggregates, even after just one hour (Figure 2A). Furthermore, the PDI in PS solutions was much higher than in Ag solutions (Figure 2B), indicating a wide distribution size. After 24 hours, the Z-average of both types of NP solutions increased in a concentration-dependent manner although overall the size of aggregates in PS solutions remained greater than those in Ag solutions (24.73 ± 9.75 vs. 2.56 ± 0.56 µm, at 1 mg/L concentration respectively). The disparity is also qualitatively reflected in the images of the NP solutions in ASW sitting idle for 24 h (Figure 2C), where increasing concentration led to an increase in the agglomeration and resulted in sedimentation at the bottom of the test tubes (markedly larger and darker for PS NPs). The difference in aggregation behavior may at least be partially attributed to the presence of citrate capping agent in Ag NPs ^39,89,90^, as opposed to PS NPs where a trace amount of surfactant is present in the as-received concentrated solution but would be diluted during suspension in ASW. Additionally, the difference in particle morphology (spherical vs. irregular, Figure 1) can affect the aggregation behavior, with predominant crystal faces ^91^ (in irregular particles) causing slower diffusion. Accordingly, whereas the PDI of PS particles decreased with increasing concentrations after 24 hours due to the development of microscale aggregates of comparable size, the PDI of Ag particles was highest at this timepoint for all concentrations reflecting nonuniform particle aggregation.

**Figure 2.**
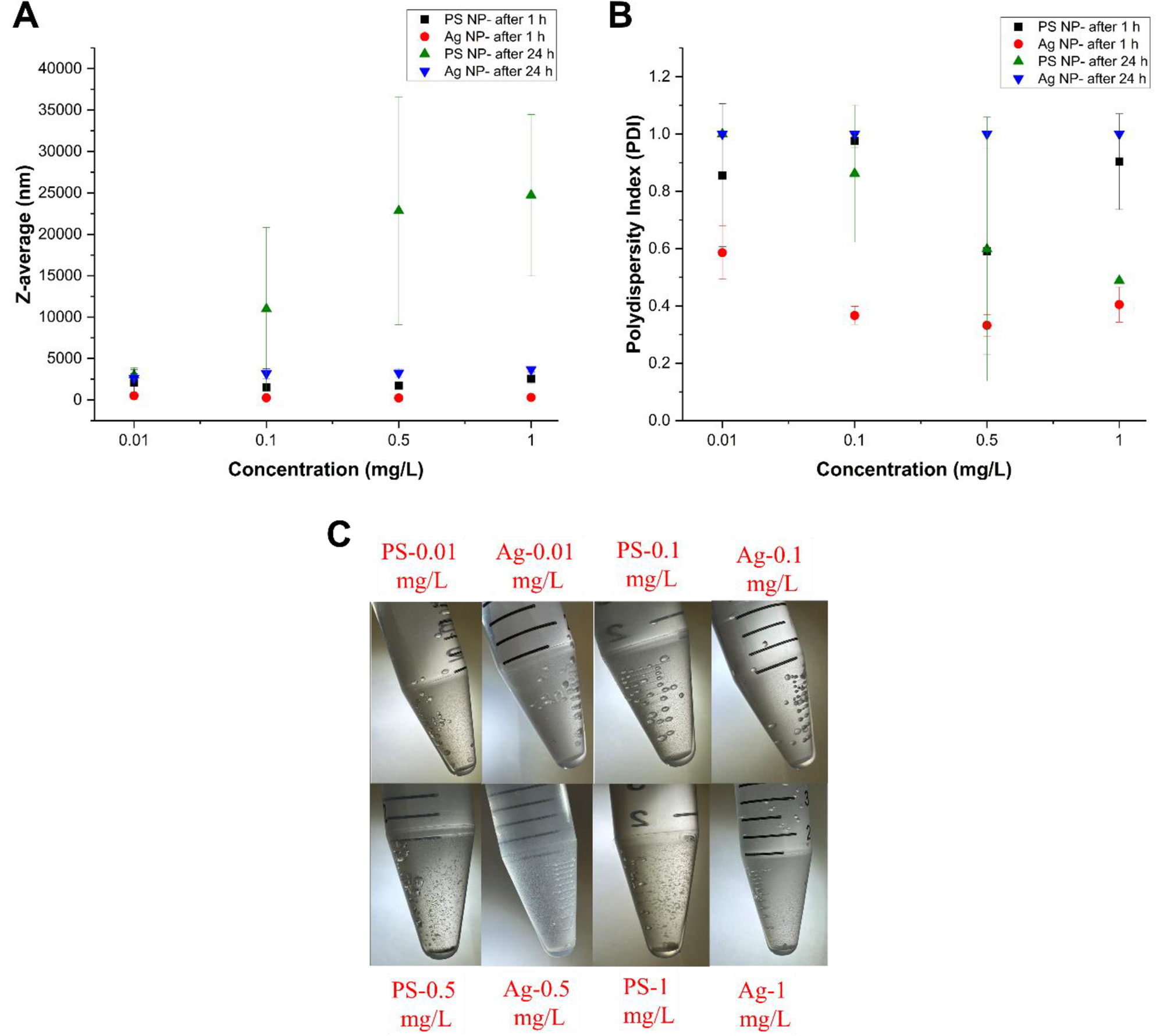
Aggregation behavior of different concentrations of polystyrene and silver nanoparticles in artificial saltwater (ASW). (A) Z-average values and (B) polydispersity index (PDI) of the NP solutions measured by DLS after 1 hour and 24 hours. Values are expressed as mean ± standard deviation (n=3 at each concentration). (C) Photos demonstrate the aggregation and subsequent sedimentation of various NP solutions in ASW after 24 hours at 25⁰C.

### 3.2 Hatching stages of *Artemia,* real-time monitoring and interaction with nanoparticles during hatching

During the oviparous method of reproduction, *Artemia* females create diapause cysts that can persist up to 28 years without detectable metabolic activity ^92–94^. When environmental conditions are favorable (temperature and salinity primarily), the cysts resume metabolic activity and are able to hatch ^95–97^. The hatching process consists of four distinct stages, encompassing the encysted state to swimming *Artemia* ^98^. Figure 3 shows the photomicrographs of the morphological changes of the hatching cysts at different stages (Figure 3A) and the corresponding oxygen consumption behavior (Figure 3B) when the cysts were not exposed to NPs (control). As well as the physical transformation, each of these hatching stages can be distinguished based on the respective energy metabolism ^98–100^. Energy metabolism is directly related to oxygen consumption, and as mentioned earlier in section 2.3, this can be measured by the on-chip oxygen sensor through depletion in the dissolved oxygen concentration (DDOC) (Figure 3B). Supplementary Figure 3 shows that compared to the DDOC in the blank experiment, the DDOC with cysts with or without NP (control) changed significantly over time, indicating that, even though the hatching chip material PDMS is oxygen permeable ^101,102^, the oxygen consumption of the cysts was significantly greater than the oxygen permeation. We note that the DDOC may be affected by the oxygen consumption of microorganisms that may be present in the nonsterile *Artemia* cysts utilized in this study; this effect was not accounted for in the current study and requires additional investigation. However, the negligible change in blank experiment over 24 hours (Supplementary Figure 3) indicates that hatching chip and ASW did not affect the DDOC, therefore indicating the absence of such potential microorganisms in both the hatching chip and ASW. Figure 3B also depicts the average rate of oxygen consumption (aROC) for each stage, calculated using equation (1) and (3) as described in section 2.3.

**Figure 3.**
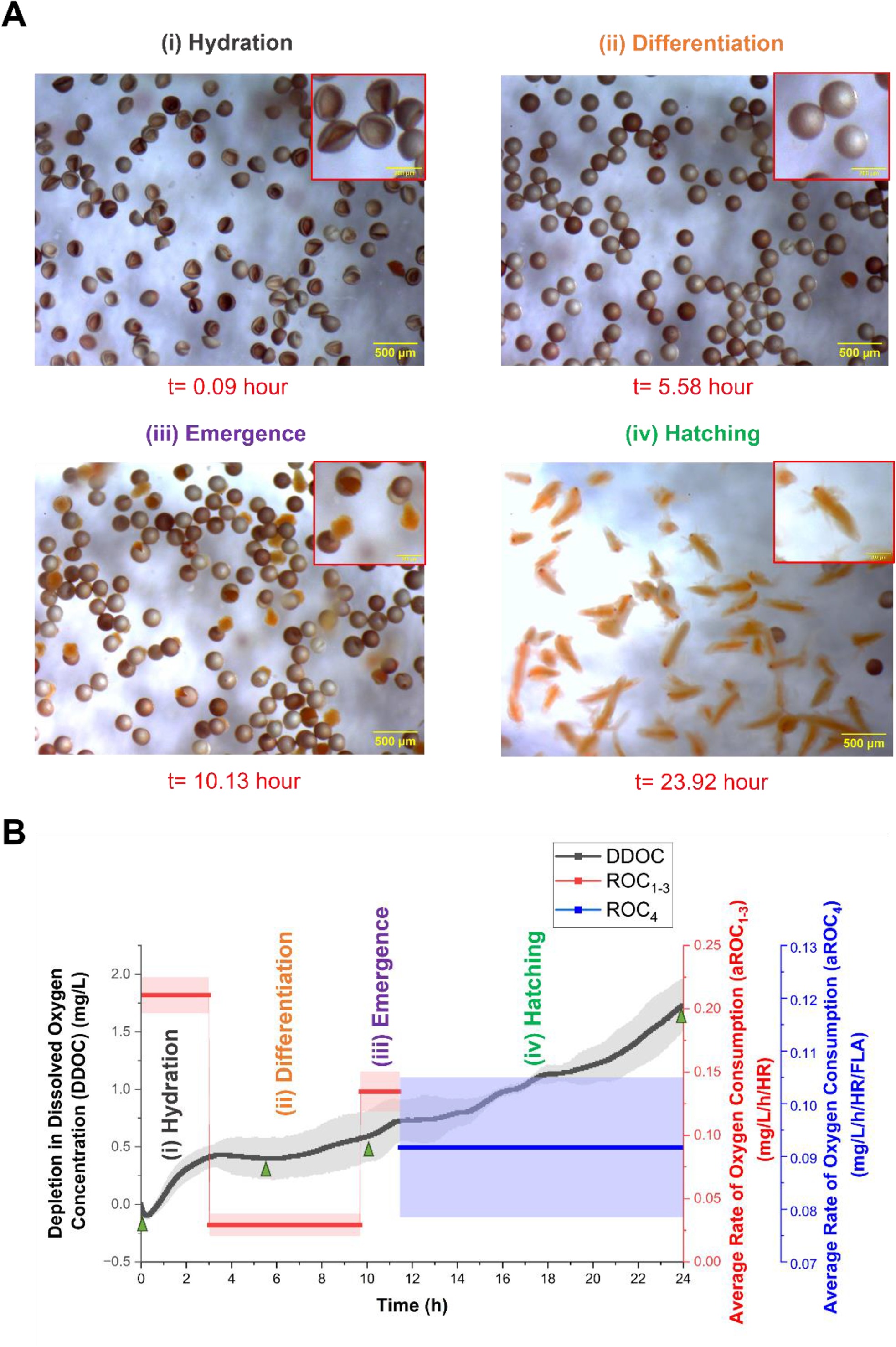
Photomicrographs and O_2_ sensor data of the four stages of hatching process. (A) Photomicrographs of the four stages of hatching captured with an optical stereomicroscope [the inset depicts the morphology of the cysts and nauplii at various stages (scale bar= 200 µm)] (scale bar = 500 µm). (B) On-chip detection of the change in oxygen consumption of *Artemia* cysts at the four stages of hatching measured through the depletion in dissolved oxygen concentration (DDOC) and average rate of oxygen consumption (aROC) when exposed to no NPs (control) (The bold lines in the graph indicate the average, and shaded region indicates the standard deviation (n≤3). Red line/shaded region corresponds to the aROC_1-3_ axis and blue line/shaded region corresponds to the aROC_4_ axis on the right Y-axis. The green triangles in Figure 3B depict the DDOC at the time points corresponding to the photomicrographs in Figure 3A).

The initial step of hatching is the hydration stage, which occurs when a dormant cyst is immersed in saltwater at favorable temperature and salinity levels ^57^. The cysts absorb water through osmosis and inflate, transforming from cup-shaped to nearly spherical [Figure 3A(i)] ^103^. As the cysts imbibe water, cellular energy metabolism is reactivated along with RNA and protein synthesis capacity modifications ^99,104^ resulting in an increase in aROC [Figure 3B(i)] ^57^. The second stage is differentiation, during which the spherical shape of the hatching cysts remains unchanged [Figure 3A(ii)] and there is no internal cell division or DNA replication ^99,104^. Accordingly, the oxygen demand remains unchanged throughout this stage resulting in a lower aROC [Figure 3B(ii)] ^57^. During the third stage of emergence, the *Artemia* embryos emerge from the cysts, which rupture as a result of increasing internal turgor pressure caused by the synthesis of glycerol from trehalose ^100^; a sugar produced and retained throughout the early stages of embryo development, from fertilization until the start of dormancy ^105^. As embryos emerge from rupturing cysts [Figure 3A (iii)], they require more energy, resulting in an increased aROC [Figure 3B (iii)] ^57^. The final stage of this process is hatching, when the embryos leave the cyst shells within a hatching membrane and start swimming [Figure 3A(iv)]. At this stage, the embryos expend energy for swimming activity, but the combined oxygen requirement is lower than that required to break the cyst corresponding with a reduced aROC compared to emergence [Figure 3B(iv)] ^57^.

As the cysts hatch through the aforementioned stages, it is crucial to understand how they interact with NPs, if present. NPs might penetrate through the fracture of the cyst shell during hatching and the inner 3D porous structure (Figure 1), which might interfere with the hatching process. The morphology of the cysts hatched in the presence of NPs and the distribution of Ag NPs on the cysts were investigated using SEM (Figure 4A) and EDS (Figure 4B), respectively. For this, as previously stated in section 2.2, *Artemia* cysts were hatched in PS and Ag NP solution in DIW to avoid obscurity of the cyst morphology due to salt crystal formation from ASW evaporation (Supplementary Figure 2A). Both the PS and Ag NPs were observed to aggregate in the saltwater solution (Figure 2), and it is of value to understand how aggregates of NPs interact with the cysts during hatching. To simulate this, as previously mentioned in section 2.2, *Artemia* cysts were also hatched in solution of bigger PS (2, 10 µm) and Ag (100 nm) NPs in DIW and their morphology was observed under SEM (Supplementary Figure 2). From the SEM images, the cysts were observed to be penetrated by NPs through the cracks that were formed during the process of cyst expansion/emergence caused by hydration and an increase in internal turgor pressure. The inner 3D honeycomb-like structure of the cysts also facilitated this penetration. This was confirmed by non-visible inner pores of the cysts that were hatched in the presence of NPs [Figure 4A (ii and iii) and Supplementary Figure 2B] compared to the clearly visible pores in the cyst shells that were hatched in only DIW [Figure 4A(i) and Supplementary Figure 2B(i)]. Furthermore, the cysts were observed to be covered by NPs (Supplementary Figure 2C) due to the possible physical attraction between the NPs and various functional groups on the surface of the cysts ^106^, such as amide, carboxylic, and so on. EDS analysis also revealed that the Ag NPs were distributed on the surface of the inner porous structure of cysts (Figure 4B). In addition, the SEM images demonstrate a reduction in the infiltration capacity of PS NPs into the 3D inner porous structure with smaller pore size (maximum of 900 nm) as the size of PS NPs increase (Supplementary Figure 2B).

**Figure 4.**
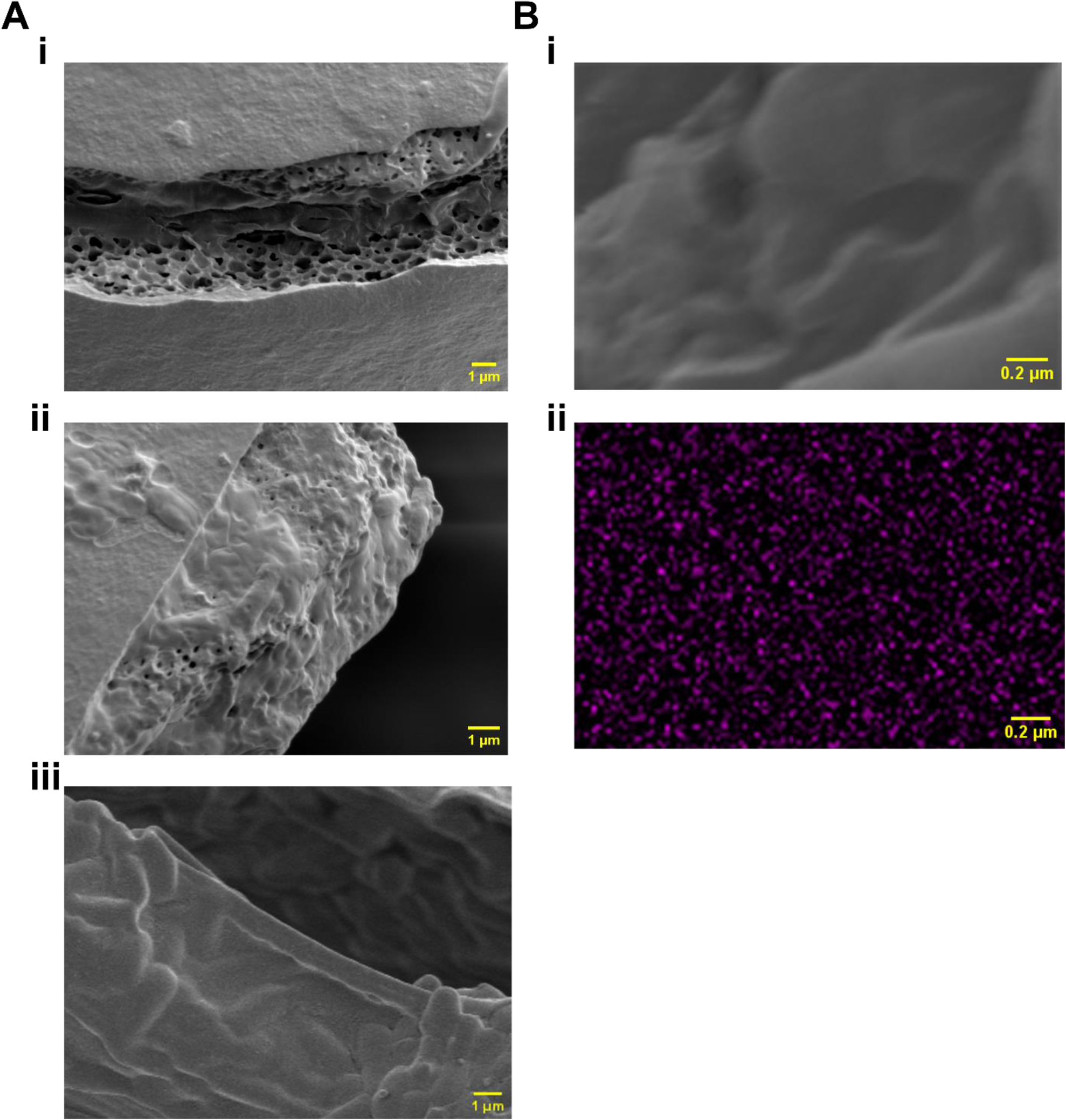
Scanning electron microscopy (SEM) images and Energy dispersive X-ray spectrometry (EDS) analysis of the cysts hatched in solution of PS (50 nm size) and Ag (40 nm size) NPs in deionized water (DIW). (A) SEM images of cyst shell crack when cysts were hatched in- i) only DIW ii) PS NP and iii) Ag NP (scale bar = 1 µm). (B) EDS analysis shows- i) the inner porous structure of cyst, ii) distribution of Ag NPs on the same surface as B(i) (scale bar = 0.2 µm).

### 3.3 Effect of PS and Ag nanoparticles on the duration of different hatching stages and hatching rate

Previously, we demonstrated that the duration of the hatching stage can be correlated with hatching survival ^57^. Figure 5A illustrates the effects of PS and Ag NP exposures on the duration of each of the four stages of hatching. Overall, the effects of the presence of either PS or Ag NPs on hatching duration and oxygen consumption are non-monotonic in terms of particle concentration. This finding is in agreement with literature that suggests toxicity is influenced by particle size ^107^, as well as the observed patterns in changes in aggregate size over time in our study (Figure 2), which were influenced by particle type and concentration.

**Figure 5.**
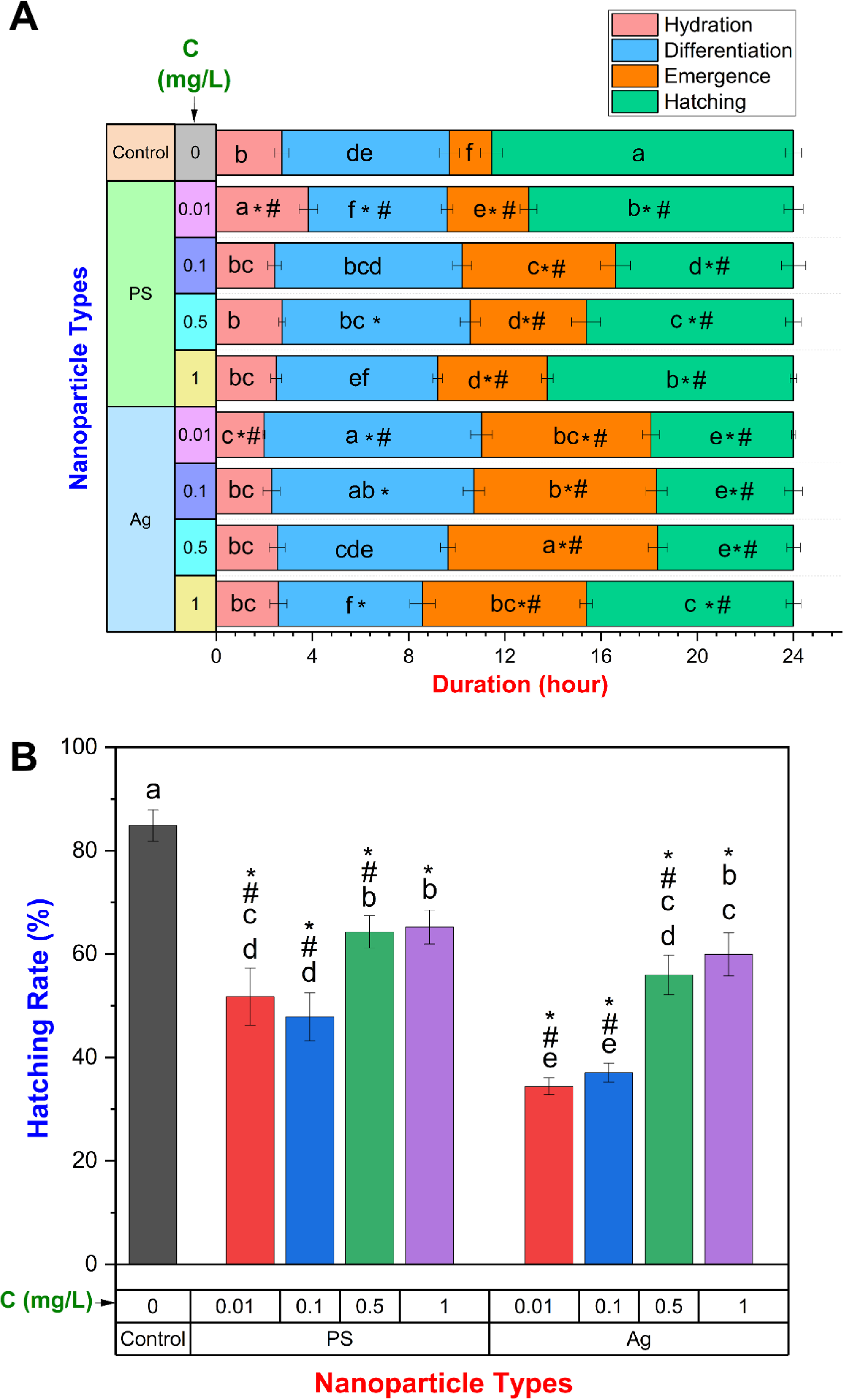
Effect of PS and Ag NPs of varying concentrations on- (A) the duration of different stages of hatching of *Artemia*, and (B) hatching rate. Values are expressed as mean ± standard deviation (n≤3). Means that do not share the same letter are statistically significantly different (*p*<0.05) [in each stage in Figure 5(A)]. * and # are indicated separately to emphasize data that differs significantly from control (without NP) and hatching under two different NPs treatments (PS and Ag) at the same concentration, respectively) [in each stage in Figure 5(A)]. “C” represents concentrations of the NP treatments.

Beginning with hydration, we find that for both types of NPs, the hydration duration affected significantly (Figure 5A-red bars) only at the lowest concentration tested (0.01 mg/L). At low concentrations, the aggregate size is smaller (Figure 2) allowing the NPs to permeate the fracture generated during hydration (Figure 4), which would alter the osmotic gradient and inhibit the cysts ability to imbibe water. At the same time, the absence of an effect on hydration duration at higher concentrations suggests that the adhesion of NPs to the exterior surface of the cysts does not affect the hydration process. The difference in the hydration duration in the presence of the two types of NPs may be attributable to differential toxicity (discussed later in section 3.5).

The differentiation stage (Figure 5A-blue bars) is significantly affected by both the type and concentration of nanoparticle in different ways. In the presence of PS NPs, the duration decreased at the lowest tested concentration, followed by an increase with rising NP concentrations up to 0.5 mg/L, and subsequent decrease. In contrast, in the presence of Ag NPs, the differentiation duration increased at the lowest measured concentration, but decreased with higher concentrations. During the differentiation stage the cysts progress towards the emergence stage when the shell starts to fracture due to the rising turgor pressure. The fracturing cysts may be penetrated at this time by the NPs more readily (Figure 4) and induce toxicity. The NPs could possibly interfere with cellular differentiative processes and cause genetic damage ^108^, and result in a longer differentiation stage as compared to the control. The reduced effect with increasing concentration is likely due to lower bioavailability due to sedimentation. A similar reduction in bioavailability with increased aggregation of NPs in saltwater was reported in previous studies, such as with TiO_2_ NPs ^109^, and cerium oxide NPs ^110^.

The presence of either NP significantly lengthened the duration of the emergence stage in comparison to the control at all concentrations tested (Figure 5A-orange bars). One potential explanation is that NPs may result in a deficit of bicarbonate ions which can influence transport across the cytoplasmic membrane to reduce osmotic potential ^111^ in which case the cyst shell breaks, but the inner cuticular membrane retains its integrity owing to its elasticity, which can hinder emergence and lengthen the process ^111^. The increased duration in emergence was significantly greater in the presence of Ag NPs at all concentrations when compared to PS NPs. The observed phenomenon may be associated with the elevated toxicity of metals, as previously observed in the presence of mercury ^111^, cadmium ^112^, zinc ^112,113^, iron ^113^, lead ^113^, and nickel ^113^ which has been attributed to the increased hindrance to ion transport in the membranes of salt glands, responsible for producing osmotic potential ^113,114^.

Finally, the hatching stage duration is the remaining time after the first three stages within the 24 hours of the experiment (Figure 5A-green bars). The presence of both NPs not only markedly altered the durations of the hydration, differentiation, and emergence stages, but affected the remaining time for the hatching stage within the experimental timeframe. Specifically, the hatching stage was significantly shorter in the presence of Ag NPs compared to PS NPs, as well as the control.

The hatching rate (HR) of *Artemia* cysts in the presence of various concentrations of PS and Ag NPs after 24 hours is presented in Figure 5B, calculated using equation (2). Similar to the duration, the presence of both types of NPs significantly inhibited the hatchability of *Artemia*. However, in agreement with the previous observations in duration (Figure 5A), the negative impacts decreased with increased NP concentration, resulting in an increased HR. Overall, Ag NPs inhibited *Artemia* hatching significantly more than PS NPs with the exception of the highest concentration of 1mg/L, at which the hatching rate did not differ. The lowest hatchability (34.42 ± 1.66%) among the tested conditions was observed in the presence of Ag NPs at a concentration of 0.01 mg/L, which was 59.44% lower than the control. Effects on duration of different stages of hatching in the presence of NPs (Figure 5A), resulted in the inhibition of *Artemia* hatching, especially in the presence of Ag NPs, which affected the emergence and hatching stages significantly more than PS NPs. Specifically, interference with cellular differentiative process ^108^, reactive oxygen species (ROS) formation ^115^, damage in DNA moieties ^116^, cuticle damage ^117^, and perturbed osmotic gradient ^113,114^ may have contributed to the low hatching rate.

### 3.4 Effect of PS and Ag nanoparticles on oxygen consumption

Average rate of oxygen consumption at the hydration stage (Figure 6A) reveals that the oxygen consumption significantly increased in the presence of Ag NPs across all concentrations, except for 0.5 mg/L, when compared to control. However, in the presence of PS NPs, the oxygen consumption did not change significantly except at 0.1 mg/L concentration. Although, NPs did not significantly affect the hydration duration (except at 0.01 mg/L) (Figure 5A-red bars), the observed increased oxygen demand could be attributed to increased metabolic cost incurred due to the adherence of the NPs on the cyst surface (Supplementary Figure 2) and resulting hindrance to the resumption of metabolism in dormant cysts. During the differentiation stage, oxygen consumption tended to remain constant (Figure 6B), which was also observed in our previous study ^57^, and the duration increased in the presence of NPs (Figure 5A-blue bars), but there was no significant change in aROC compared to control. Consistent with the delayed emergence stage observed in the presence of NPs (Figure 5A-orange bars), the aROC at the emergence stage increased significantly when PS NP concentration increased to 0.1 mg/L, with no further significant change when concentration increased further (Figure 6C). Similarly, the aROC increased when the concentration of Ag NPs increased to 0.1 mg/L and then declined with increasing concentration. A significant increase in the aROC was observed during the hatching stage (aROC_4_) upon increasing the concentration of PS NP to 1 mg/L. Similarly, an increase in aROC was observed when Ag NP concentration increased beyond 0.1 mg/L, except at 0.5 mg/L. A similar increase in oxygen consumption has been observed in other aquatic species exposed to different NPs, e.g., *Perca fluviatilis* exposed to AgNO_3_ ^118^, *Brachidontes pharaonic* exposed to Ag NPs ^119^, *Fundulus heteroclitus* exposed to Cu NPs ^120^ and *Apistogramma agassizii* and *Paracheirodon axelrodi* exposed to Cu and CuO NPs ^121^. This effect could be attributed to gill impairment, characterized by hypertrophy and hyperplasia, as well as the proliferation of mitochondrion-rich cells (MRC) caused by the buildup of nanoparticles (NPs) ^121^. Damage to gill structure can lead to compromised respiratory gas exchange, increased energy demand for osmoregulatory processes, and subsequent increase in oxygen consumption ^121,122^.

**Figure 6.**
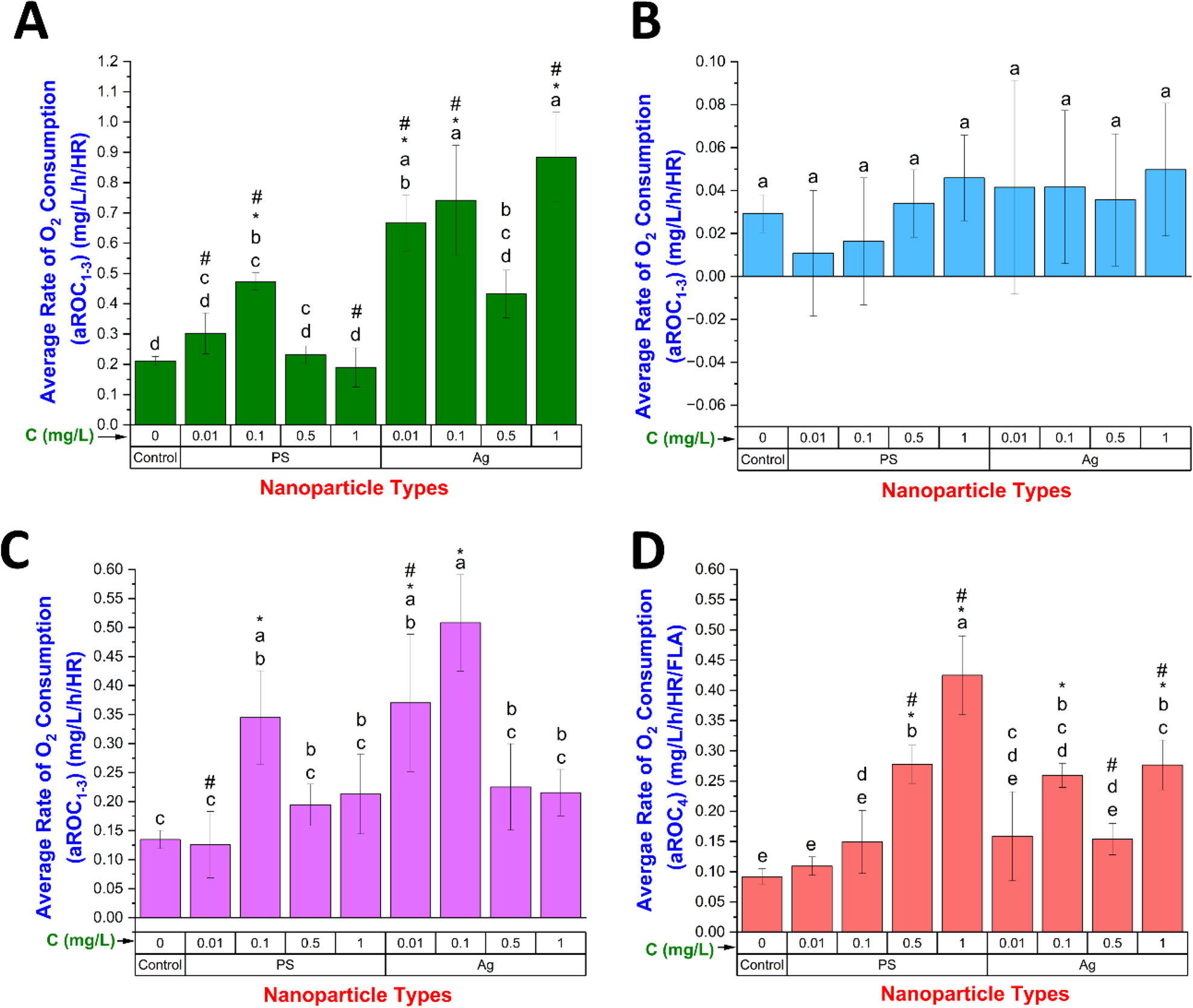
Effect of PS and Ag NPs of varying concentrations on the oxygen consumption of *Artemia*. (A-C) Average rate of oxygen consumption (aROC) during the first three pre-hatching stages (hydration, differentiation, and emergence) normalized based on duration of stages, and hatching rate (HR). (B) aROC at the hatching stage normalized based on duration of stage, HR and fraction of live *Artemia* (FLA). Values are expressed as mean ± standard deviation (n≤3). In each category of data, means that do not share the same letter are statistically distinct. “C” represents concentrations of the NP treatments.

### 3.5 Bioconcentration of PS and Ag nanoparticles, acute toxicity, and swimming speed alteration

As the mouth and anus are not yet developed, newly hatched nauplii (Instar I) are unable to consume food -or ingest nanoparticles- and depend on the yolk sac for nutrition ^123^. Nonetheless, due to the high adsorption capabilities of NPs, they may cover the body surface and gills of nauplii ^83^, negatively impacting metabolism and development. Depending on the exposure conditions, the average remaining time for the hatching stage within the experimental 24h period in this study ranged from approximately 7.74 to 12.56 hours (Figure 4). Within 6-8 hours of hatching, nauplii Instar I transform into the second larval stage (Instar II) ^124^. At the instar II stage, larvae begin to ingest food after sifting it with their antennae. *Artemia* are a nonselective filter feeders that can wconsume anything less than 50 µm in size ^53^. Figure 7 depicts fluorescence microscopy and bright field images of *Artemia* nauplii in the second instar stage hatched in varying concentrations of nanoparticles. As the Ag NPs were not fluorescent, they are imaged in bright field only. *Artemia* ingested a substantial quantity of NPs that accumulated primarily in the digestive tract (Figure 7). Figure 7A demonstrates the absence of red fluorescent PS NP in the control, whereas Figure 7B illustrates the presence of PS NPs (red particles). The blue fluorescence observed in Figure 7A, and B is indicative of the autofluorescence emitted by *Artemia*. The increase in red fluorescence intensity relative to the control reflects the concentration-dependent uptake of NPs by the nauplii (Supplementary Figure 4). Bright-field microscopy images show the digestive tract of the control group (depicted with red arrow) as nearly empty (Figure 7C), but the digestive tract of *Artemia* exposed to Ag NPs has a dark area suggesting it was filled with Ag NPs (Figure 7D). The fluorescence microscopy images indicate that the body surface of *Artemia* nauplii is also decorated by aggregates of PS NPs, which are more visible at higher concentrations, as depicted in Figure 7B(iv). However, this observation could not be validated through the bright-field microscopy method due to the non-fluorescence of Ag NPs.

**Figure 7.**
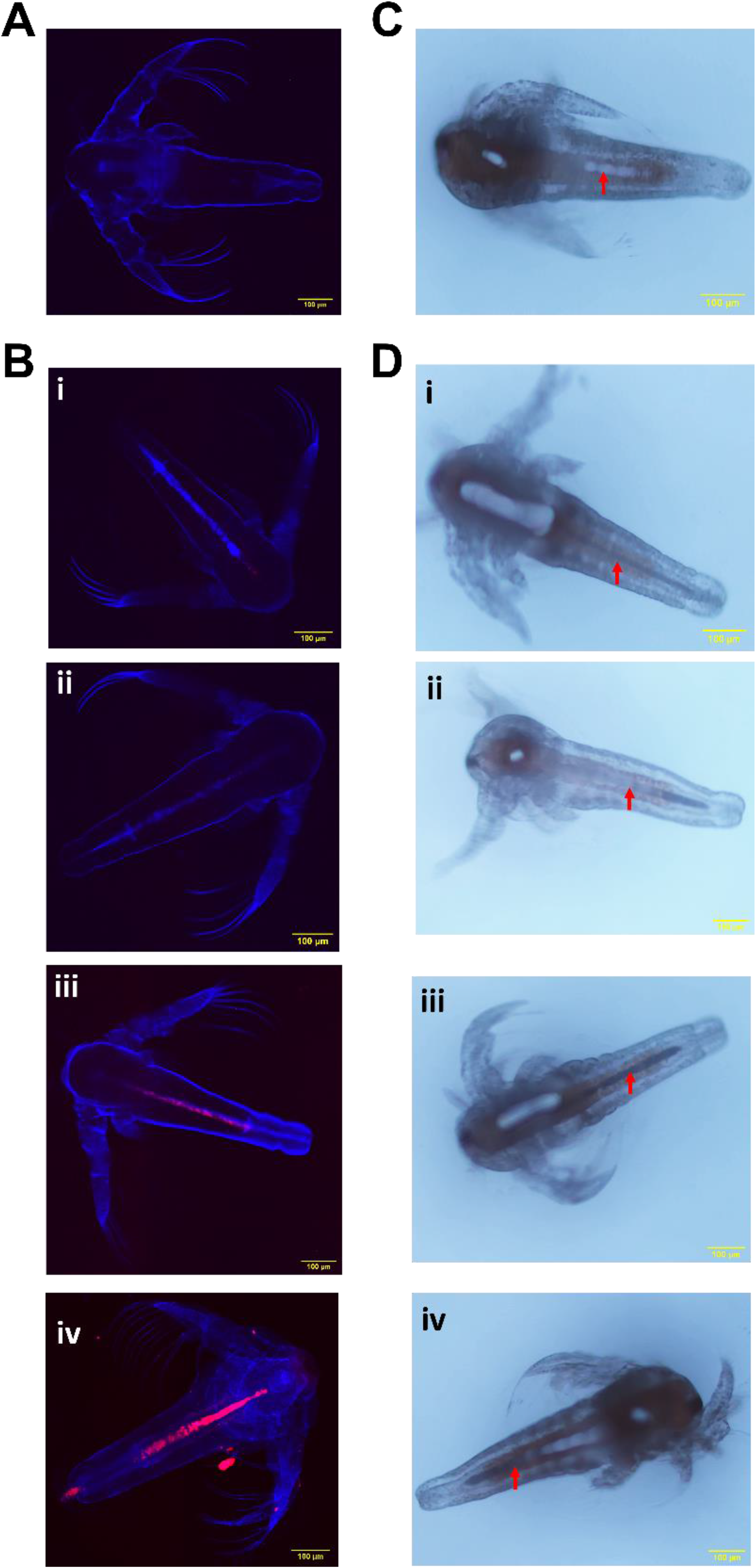
Fluorescence microscopy and bright field images of live *Artemia* exposed to different nanoparticle treatments for 24 hours during hatching. Fluorescence microscopy images of- (A) control, (B) Polystyrene (PS) exposed *Artemia.* Bright-field images of- (C) control, and (D) Silver (Ag) exposed *Artemia*. In figures B and D, i, ii, iii, and iv represent *Artemia* exposed to 0.01, 0.1, 0.5, and 1 mg/L concentrations of NPs, respectively (scale bar= 100 µm). The red arrows in Figures 7C and 7D depict the location of *Artemia*’s digestive tract.

Figure 8 depicts the mortality rate and swimming speed alteration of *Artemia* hatched in the presence of PS and Ag NPs over 24 hours, as estimated by equations (5) and (6) respectively. In comparison to the control, *Artemia* nauplii mortality was markedly elevated in the presence of both NPs. Mortality increased with rising NP concentration, while hatchability inhibition decreased as NP concentrations increased due to decreasing bioavailability (Figure 5). Although at higher concentrations the NPs aggregated more (Figure 2), after hatching is complete, free-swimming *Artemia* may be in more contact with NP aggregates, effectively increasing the bioavailability. Moreover, when they reach second instar stage, the larger aggregates of NPs (at higher concentrations) would be more readily ingested than individual NPs or smaller aggregates. At the maximum concentration of NPs studied (1 mg/L), the *Artemia* mortality rate in the presence of PS and Ag NPs was 20.48 ± 3.61% and 40.66 ± 4.48%, respectively, comparable to the results reported in the literature ^47,48,53^. This indicates that Ag NPs were more toxic to newly hatched *Artemia* than PS NPs (almost 43% higher at 1 mg/L concentration). This finding corresponds with previous studies which show poorer gut health, increased oxidative stress, and DNA damage caused by Ag compared to PS in *Drosophila* ^125,126^. In a manner consistent with observations in other species, the uptake of NPs, as observed in Figure 7, may impair the digestive tract ^50,53,127^, infiltrate body tissues ^83,128^, increase the formation of hazardous reactive oxygen species (ROS) ^50,129,130^, reduce body defense mechanisms like superoxide dismutase (SOD), catalase (CAT) ^50,51^, and cause tissue ^53^, DNA and mitochondrial damage ^53,83^, which might be attributed to the mortality of *Artemia* in the presence of NPs.

In addition to causing mortality of hatched *Artemia*, the presence of NPs significantly altered the swimming speed of *Artemia*, as determined by swimming speed alteration (SSA) using equation (5). The swimming speed of *Artemia* exhibited a decrease as the concentration of PS NP increased to 0.1 mg/L, resulting in an increase in SSA. However, a further increase in PS NP concentration resulted in a significant decrease in SSA (increase in swimming speed). Nevertheless, overall, the swimming speeds exhibited a positive value, indicating that swimming speed was significantly slower than that of the control, which was similarly seen in other marine animals exposed to PS NPs ^131–134^. In contrast, when the concentration of Ag NP increased, swimming speed increased, and SSA became more negative. Reduced swimming or hypoactivity suggests a protection reaction against environmental stressors ^135^, or could be related to several factors, such as oxidative stress ^134,136^, visual impairment ^137^, neuromotor deficits ^138^, etc. Conversely, higher swimming speed or hyperactivity may infer an escape response from a hazardous environment ^135,139^, or it could be related to psychostimulant/convulsant action or anxiety-like behavior ^138^. With greater acute toxicity (Figure 8A), Ag NP was observed to deliver more stimulation as an overcompensation for a homeostasis imbalance ^140^ and encouraged faster swimming speed as an escape response from a dangerous environment ^139^, or caused hyperactivity as observed in two other aquatic invertebrate larvae. A similar increase in swimming speed was observed when *Daphnia magna* was exposed to Ag NP ^141^. Related to this the duration of the hatching stage within the 24-hour experimental timeframe decreased significantly in the presence of Ag NPs vs PS NPs (Figure 4) such that the hatched nauplii were exposed to PS NPs for a longer period. Overall, with the alteration in swimming speed, oxygen consumption and thus energy expenditure also increased (Figure 6D), which may inhibit development, foraging, reproduction and enhance susceptibility to predators ^142^.

**Figure 8.**
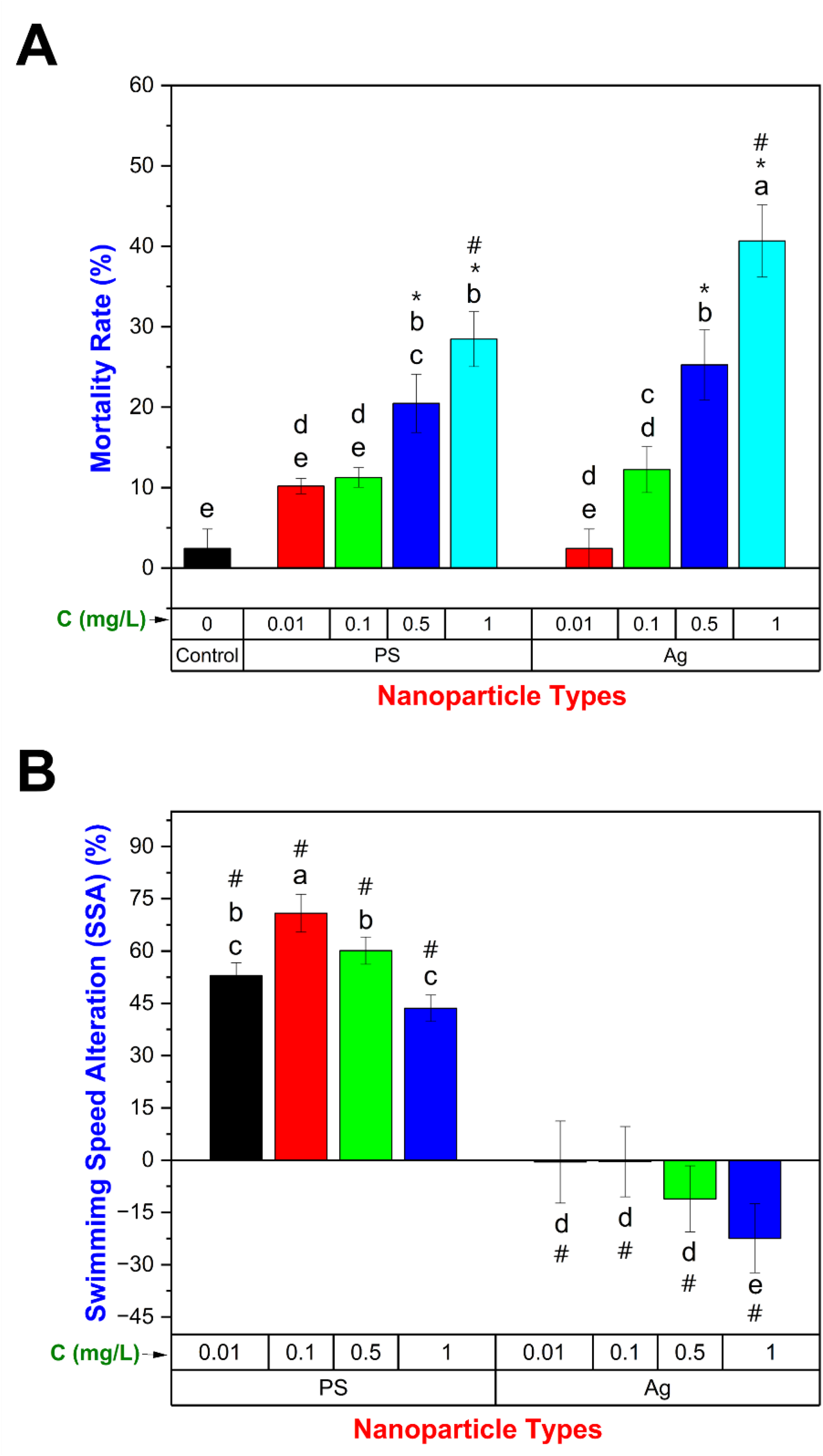
Effect of different concentrations of nanoparticles on the mortality and mobility of the hatched *Artemia* after 24 hours of hatching under various NP treatments. (A) Mortality rate, and (B) swimming speed alteration. Values are expressed as mean ± standard deviation (n≤3 at each treatment). In each category of data, means that do not share the same letter are significantly different. * and # are indicated separately to emphasize data that differs significantly from control (without NP) and hatching under two different NPs treatments (PS and Ag) at the same concentration, respectively. “C” represents concentrations of the NPs treatments.

### 3.6 Multivariate analysis

A multivariate analysis was performed to determine potential associations between the various parameters during the hatching process and nauplius stage of *Artemia* (Supplementary Figure 5) irrespective of the type of nanoparticle and concentrations. The Pearson correlation coefficient was employed to compare the parameters. Since the effects of hatching of *Artemia* cysts in the presence of NPs on the post-hatching early-stage could not be separated, as described in the methods (Section 2.7), the parameters were categorized into two groups: up to hatching (Supplementary Figure 5A) and post-hatching (Supplementary Figure 5B).

Supplementary Figure 5A shows that the duration of the hydration stage was significantly negatively associated with the durations of the differentiation and emergence stages, and positively associated with the duration of the hatching stage. In general, the durations of the differentiation and emergence stages increased in the presence of NPs at lower concentrations but decreased when the concentration increased further (Figure 4). As previously stated, this phenomenon may be ascribed to the toxic nature of the nanoparticles, and the potential reduction of toxicity resulting from aggregation and decreased bioavailability at higher concentrations. Although there was no significant difference in the hydration duration except at very low NP concentration, within this small range of hydration durations observed across the different NP exposure conditions, an increase in hydration duration was associated with a decrease in both differentiation and emergence stages durations and hence the negative correlations. As the concentration of NPs increased, the differentiation and emergence stages were observed to shorten, resulting in an increase in the remaining time for the hatching stage within the 24-hour experimental timeframe. Additionally, a negative correlation between hydration aROC and the duration of both hydration and hatching stages was observed. Conversely, a positive correlation was observed between hydration aROC and the duration of the emergence stage. aROC at the emergence stage is negatively associated with duration of hydration and hatching stages, and positively associated with the duration of differentiation and emergence stages. Furthermore, a positive correlation between the hydration stage aROC and the emergence stage aROC was found. The presence of NPs resulted in an elongation of the differentiation and emergence stages, while the hydration and hatching stages were comparatively shortened. Also, it resulted in elevated levels of oxygen consumption during the hydration, and emergence stages, suggesting an increased metabolic burden due to their toxic properties. Co-occurrence of these factors had a negative impact on the hatchability of *Artemia*. Specifically, the durations of differentiation, and emergence stages, as well as the aROC at the hydration and emergence stages-all of which were shown to be negatively correlated with the hatching rate. On the other hand, a positive relationship was observed between hatching rate and the length of the hatching stage, suggesting that a lengthier hatching stage contributed to a greater likelihood of successful hatching. In line with this, we note that in our previous study concerning the impact of temperature and salinity on the hatching process of *Artemia*, a greater hatching rate was observed when the differentiation stage was shortened and the hatching stage was extended ^57^.

Following the hatching process, *Artemia* nauplii in the early stage (nauplius stage) were impacted by the presence of nanoparticles indicated by acute toxicity and compromised swimming capabilities (Figure 8). Supplementary Figure 5B displays the Pearson correlation coefficient among factors subsequent to hatching, revealing a significant positive correlation between the hatching stage aROC and mortality rate. However, there was no significant correlation between the hatching stage aROC and hatching rate. These findings suggest that the presence of NPs induced toxicity in the newly hatched *Artemia* nauplii, leading to a reduction in the number of viable nauplii and an increase in oxygen consumption. On the other hand, swimming speed alteration (SSA) was only associated with the duration of the hatching stage (positive association). As discussed in section 3.6, positive SSA (decreased swimming speed) was seen with PS NPs when the duration of the hatching stage was comparatively longer. Conversely, in the presence of Ag NPs when the SSA was negative (increased swimming speed) a shorter hatching stage was observed.

## 4. Conclusion

In the current study, oxygen consumption was measured in real-time, and morphological changes were recorded at regular intervals to assess ***how*** different types of marine nanopollutants impact the hatching process and early stages of an aquatic organism over a wide range of environmental-relevant dosages. *Artemia*, a well-known zooplankton used in ecotoxicological studies, was the model animal, and the impacts on its hatching process, hatching rate, mortality rate, and swimming speed were evaluated. In agreement with previous studies, the presence of both NPs had detrimental impacts on the hatching performance ^53–56^. However, whereas these previous studies were confined to an analysis of the hatching rate at the endpoint, in our study, a correlation between the progression of the hatching stages and the hatching rate was demonstrated. For example, a higher hatchability was observed when the dormant cysts expended less metabolic energy (lower oxygen consumption) in the hydration and emergence stages, required less time for the differentiation and emergence stages, and had more time available for the hatching stage. Throughout different stages of the hatching process and at the endpoint (hatching rate), a trend of diminishing toxicity with increasing dosage was observed. However, the mortality rate increased dose-dependently, which is consistent with earlier studies ^47,48,51,53,56^. The observed phenomenon can be attributed to the increased aggregation, which leads to a decrease in the bioavailability of the NPs during the hatching process. However, the bioavailability of NPs increases subsequent to the completion of hatching, when the free-swimming *Artemia* nauplii contact the aggregated NPs and ingest them. The hatchability and mortality rates were significantly affected by Ag NPs compared to PS NPs across all concentrations tested, which could be attributed to increased toxicity of heavy metals ^143,144^. In addition, while swimming speed increased in the presence of Ag NPs, it decreased in the presence of PS NPs, possibly as a result of different coping mechanisms/effects in response to diverse environmental stimuli.

We anticipate that the controlled microfluidic environment developed here, coupled with integrated real-time sensing and microscopy could be used to advance the comprehensive assessment of a broad range of marine nanopollutants (nanoparticles, pesticides, fertilizers, detergents, sewage, industrial effluents, etc.), on biological processes and marine organisms ranging from zooplankton to fish larvae.

## CRediT authorship contribution statement

**Preyojon Dey:** Conceptualization, Methodology, Investigation, Formal analysis, Visualization, Writing - Original Draft, Writing - Review & Editing **Terence M. Bradley:** Conceptualization, Supervision, Formal analysis, Writing - Review & Editing, Funding acquisition **Alicia Boymelgreen:** Conceptualization, Supervision, Formal analysis, Writing - Review & Editing, Funding acquisition.

## Acknowledgments

This work is supported by the National Science Foundation (award number: 2038484, year: 2020). Graphs were plotted using OriginPro 2023 (https://www.originlab.com/). Schematics were created using Biorender (https://biorender.com/).

