## Supplementary information for "Real-time assessment of the impacts of polystyrene and silver nanoparticles on hatching process and early-stage development of *Artemia* using a microfluidic platform"

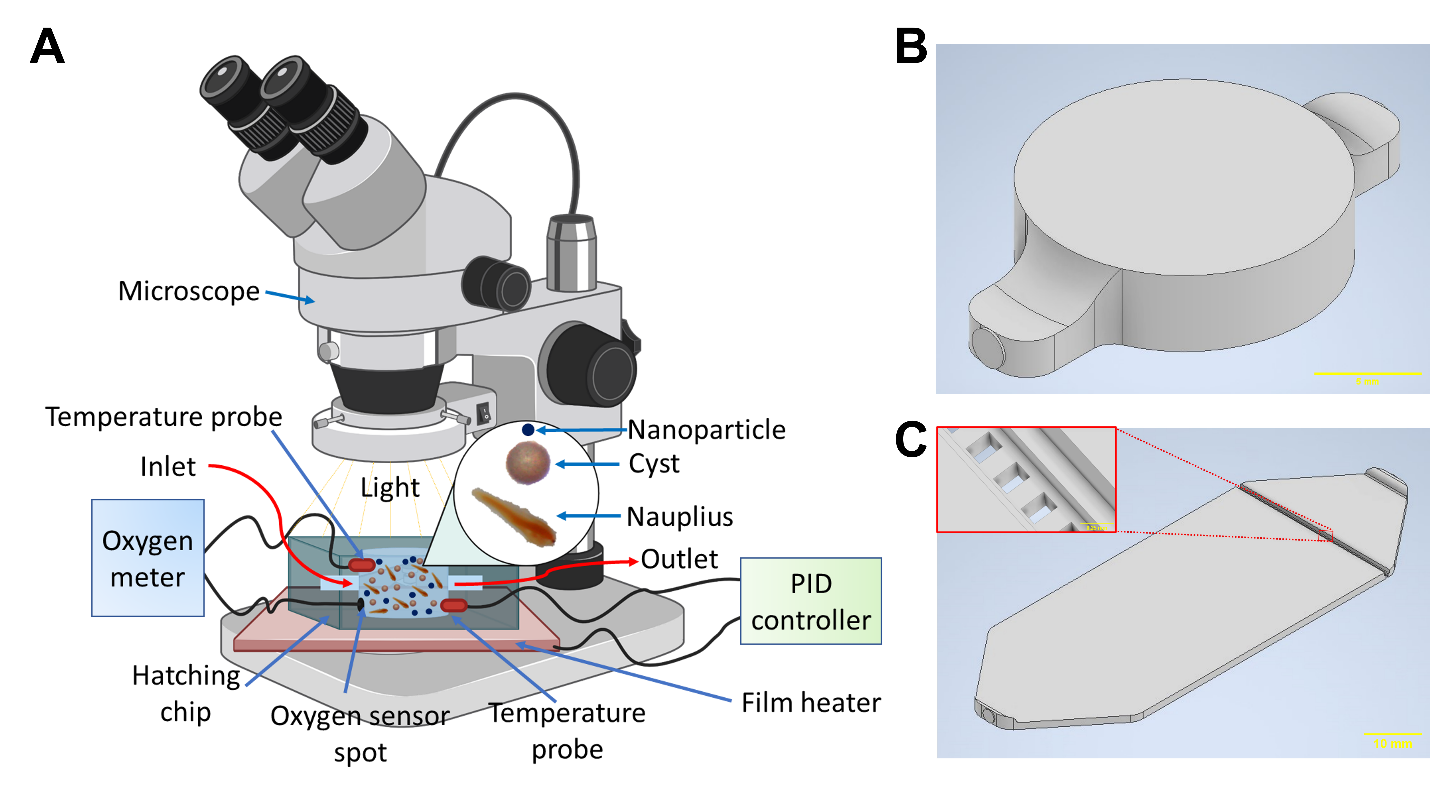

**Supplementary Figure 1.** A) Schematic diagram of the experimental setup, B) computer aided design (CAD) model of hatching chip (scale bar = 5 mm), and C) CAD model of counting chip (scale bar = 10 mm) [inset shows the micropillar structure to restrict escape of cysts and nauplii (scale bar = 0.25 mm)].

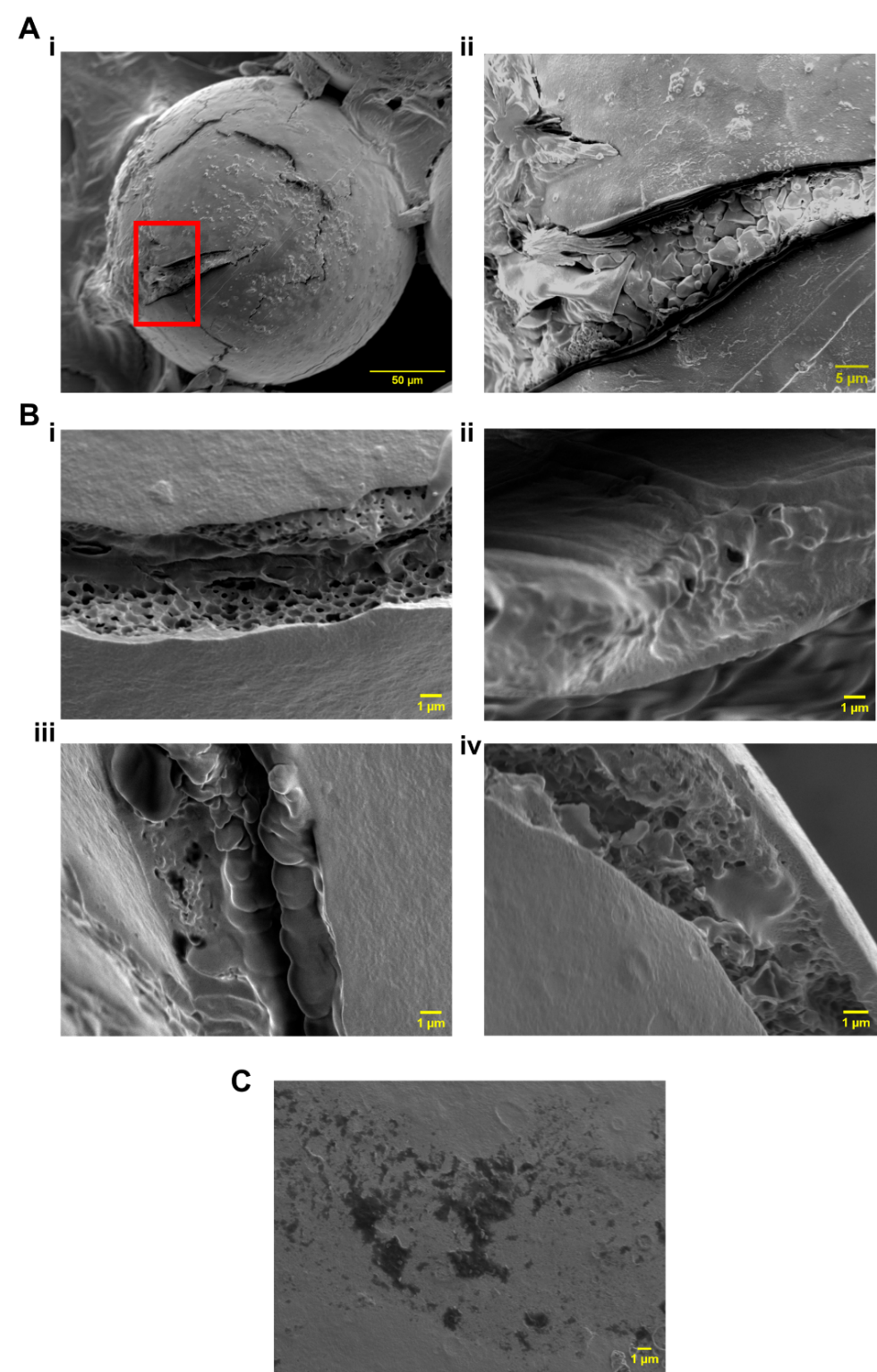

**Supplementary Figure 2.** Scanning electron microscopy (SEM) images of the *Artemia* cysts that hatched under various conditions. (A) SEM images of the cyst hatched in ASW reveal salt particles/crystals after water evaporation, obscuring the morphology of the cyst's inner 3D porous structure. (ii) is an enlarged view of the cyst shell fracture depicted in (i) (region marked with a red rectangle). (B) SEM images of the cyst shell fracture when the cysts were hatched in i) only DIW, ii) 2 µm PS NPs, iii) 10 µm PS NPs, and iv) 100 nm Ag NPs. (C) The SEM image reveals that larger PS NPs (10 µm) have adhered to the cyst surface (black area).

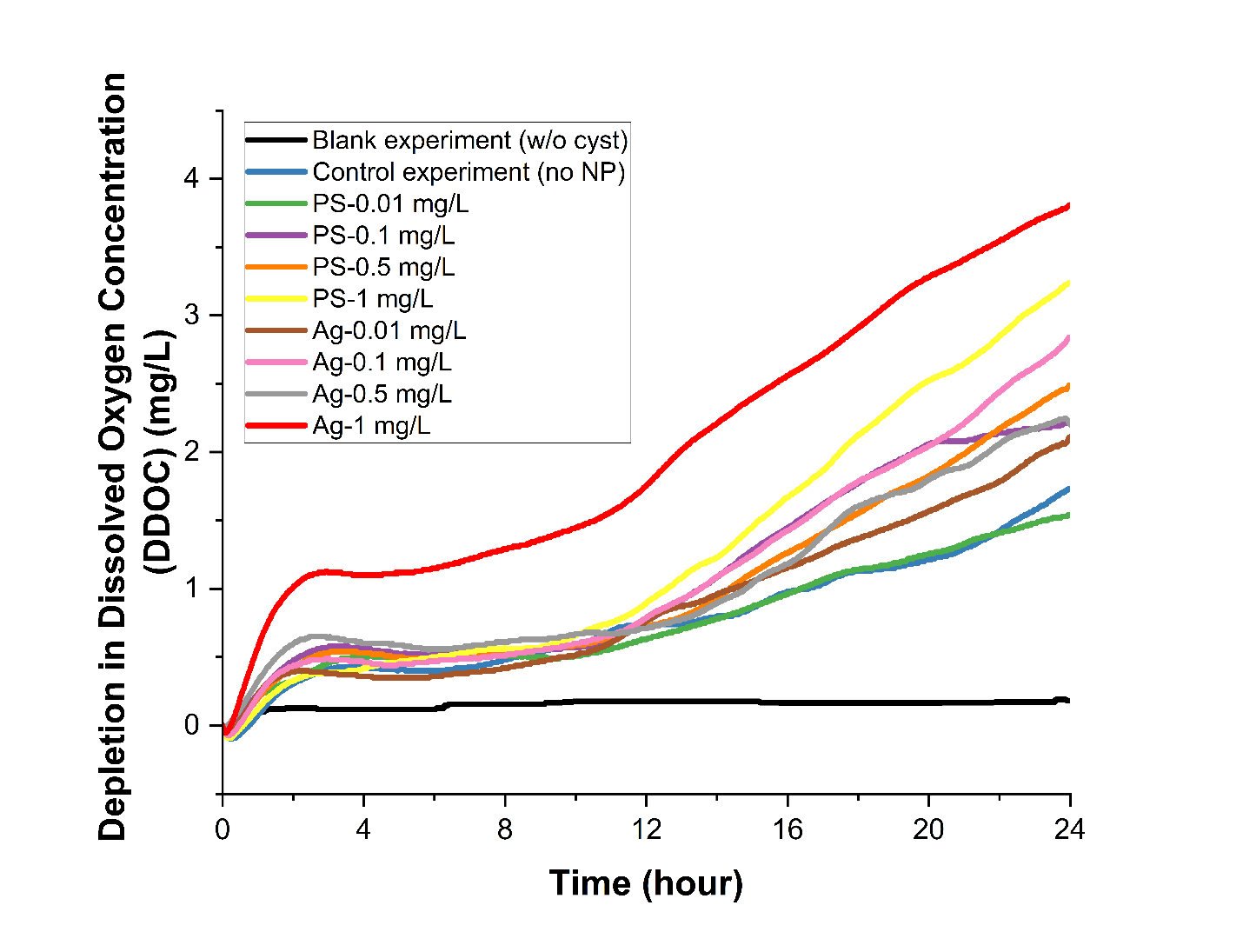

**Supplementary Figure 3.** Average depletion of dissolved oxygen concentration (DDOC) of the hatching media with time when there were no *Artemia* cysts but only artificial saltwater (blank), *Artemia* cyst were hatched without presence of any NP (control), and *Artemia* cysts were hatched in the presence of PS and Ag NPs at various concentrations (n ≥ 3).

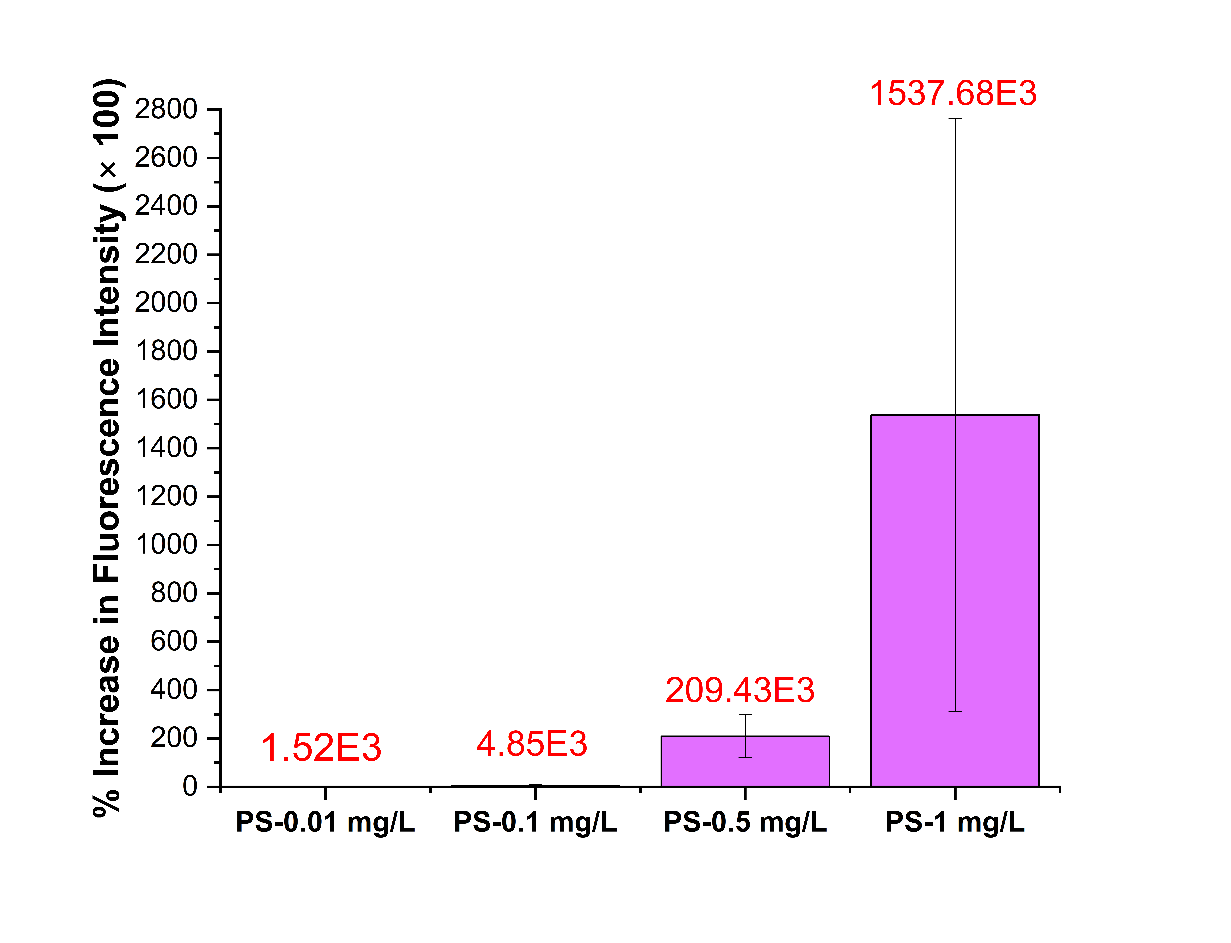

**Supplementary Figure 4.** Percentage increase in red fluorescence intensity in *Artemia* nauplii exposed to red fluorescent PS NPs as compared to control (the numbers in red color show the average % increase in fluorescence intensity) (n=3).

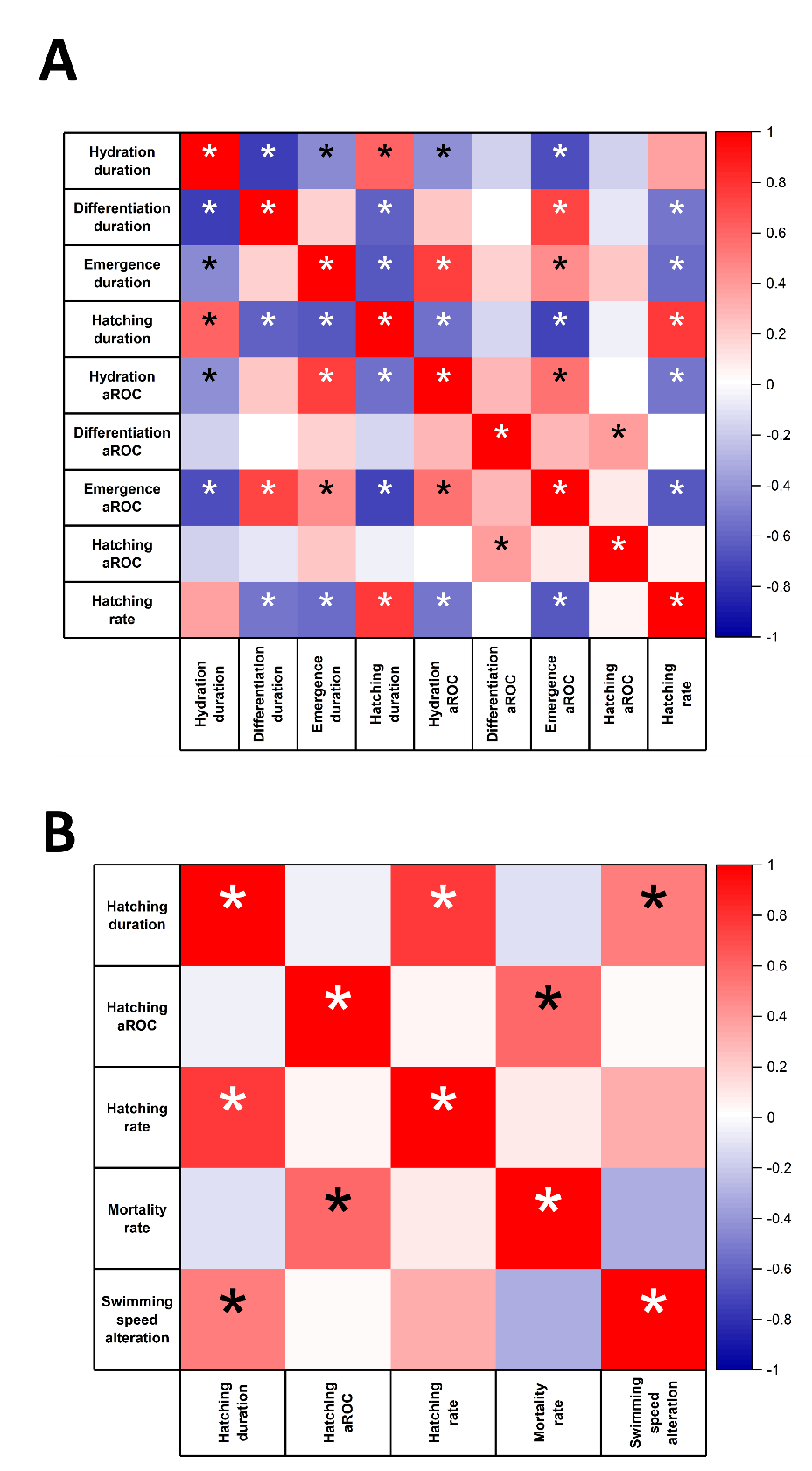

**Supplementary Figure 5.** Pearson correlation matrix on the basis of possible association among various factors of hatching process and early stage of *Artemia* analyzed in this study irrespective of NP type and concentrations (n≥27). A) Association among different factors until hatching, and B) correlation among different factors at the final stage (hatching stage) and end-point results of mortality and swimming speed alteration. The value of Pearson correlation coefficients is shown in the adjacent color scale (-1 to 1 indicating a strong negative to strong positive association). [* represents relation is statistically significant (*p*<0.05)].
